# A Flexible Transgene Integration ‘Landing-Pad’ Toolkit in Human Induced Pluripotent Stem Cells Enables Facile Cellular Engineering, Gene Zygosity Control, and Parallel Transgene Integration

**DOI:** 10.1101/2023.03.03.531057

**Authors:** Aaron H. Rosenstein, Maria Nguyen, Rasha Al-Attar, Danielle Serra, Nitya Gulati, Ting Yin, Penney M. Gilbert, Michael Garton

## Abstract

Development of a repeatable method for delivering transgene payloads to human induced pluripotent stem cells (hiPSCs) without risking unintended off-target effects is not fully realized. Yet, such methods are indispensable to fully unlocking the potential for applying synthetic biological approaches to regenerative medicine, delivering quantum impacts to cell-based therapeutics development. Here we present a toolkit for engineering hiPSCs centred on the development of two core ‘landing-pad’ cell-lines, facilitating rapid high-efficiency delivery of transgenes to the *AAVS1* safe-harbour locus using the Bxb1 large-serine recombinase. We developed two landing-pad cell lines expressing green and red fluorescent reporters respectively, both retaining stemness whilst fully capable of differentiation into all three germ layers. A fully selected hiPSC population can be isolated within 1-2 weeks after landing-pad recombinase-mediated cassette exchange. We demonstrate the capability for investigator-controlled homozygous or heterozygous transgene configurations in these cells. As such, the toolkit of vectors and protocols associated with this landing-pad hiPSC system has the potential to accelerate engineering workflows for researchers in a variety of disciplines.

## Introduction

With an ever-broadening definition, synthetic biology has the potential to deliver paradigm-shifting breakthroughs in many fields of biology, with the fields of stem cell-therapeutics and regenerative medicine being no exception. A core concept of synthetic biology is the adaptation of engineering-based approaches such as rapid-prototyping. Yet, widespread tools and platforms for simple, standardized, and rapid testing of novel mammalian gene circuits in stem-cells have yet to permeate the field and this is a major barrier for full integration of a synthetic-biological philosophy to cellular engineering. Herein, we describe and validate a flexible toolkit for repeatable stem-cell engineering, building on current progress to yield an accessible testbed for investigators to fully leverage synthetic-biological approaches for next-generation cell-based therapeutics development.

The past decade has seen the introduction of a variety of genetic engineering tools for custom cell-line generation and experimentation in stem cells. While lentiviral and transposase-based tools such as *piggyBac* are efficient means of stably transfecting hiPSCs, their target-agnosticism and variable transgene integration frequencies yield unrepeatable results^1,2^. Furthermore, positional effects from random integration events may result in transgene silencing in hiPSCs^3–5^. CRISPR-Cas9, conversely, is a highly targeted approach to stable transgene integration via homology-directed repair (HDR) at defined genomic loci in hiPSCs, but can be cytotoxic and exhibit reduced efficiency in comparison to target-agnostic methods like *piggyBac*^2,6,7^. As such, investigators have developed a compromise in the form of ‘landing-pad’ cells, pre-engineered with an exogenously-derived recombination site at an investigator-defined locus^8–13^. Delivery of transgenes is achievable by means of a site-specific recombinase. We developed a novel toolkit for gene delivery into landing-pad hiPSC cell lines and develop methodologies for straightforward stem cell engineering.

The core assumption of utilizing landing-pad cells is that transgenesis is equal to or more efficient than direct CRISPR-Cas9-based transgene knock-in, less prone to unintended mutagenesis, less toxic, and can provide predictable stable transgene expression. Safe harbour sites such as *AAVS1, ROSA26, CLYBL*, and *SHS231* have been previously characterized to provide long-term stable transgene expression, and are common landing-pad genomic insertion destinations^8,11,14,15^. Various mechanisms for effecting gene transfer into landing-pad cells have been characterized, leveraging Cre, Flp, ϕC31, and Bxb1-based recombination machinery^8–13,16^. While Cre and Flp recombinases are bidirectional, catalyzing both transgene insertion and excision, ϕC31 and BxB1 are both large serine recombinases (LSRs) and are unidirectional in the absence of a specific recombination-directionality factor^17,18^. Thus, LSRs are more practical for recombinase-based genomic integration systems such as landing-pads^19^.

Until recently, Bxb1 was the most efficient and accurate of the collection of well-characterized LSRs, while discovery of alternative LSRs such as Pa01 and BceINTa^20–22^. In the context of landing-pad cells, Bxb1 catalyzes a site-specific unidirectional recombination reaction between *attP* and *attB* DNA sequences for exchange of genetic material between the genome and a transgene delivery vector, yielding sites *attR* and *attL*, which are biologically inactive. While Bxb1 *attP* and *attB* share only partial sequence homology, matching dinucleotide core sequences (DCSs) are required for productive recombination^23^. As such, recombinase-mediated cassette-exchange (RMCE) systems comprising two *attP*/*attB* site pairs harbouring distinct DCSs can effect orthogonal recombination events, enabling transgene integration without bacterial plasmid backbone insertions^13^. Furthermore, by replacing the wild-type ‘GT’ DCS with ‘GA’, efficiency of recombination was improved^24^. More complex recombination systems have also been devised such as ‘reiterative recombination’, permitting successive integration of transgenes whereby each insertion event recapitulates a new recombination site for further engineering, and can theoretically be propagated *ad infinitum*^25^. Such a strategy can permit complex multi-expression-construct engineering in mammalian systems.

Stem cell engineering presents significant potential for both therapeutic development and basic research, providing faithful models for disease characterization and treatment. Previous recombinase-based landing-pad targeting strategies in hiPSCs such as Big-IN, STRAIGHT-IN, dual-integrase cassette exchange (DICE) system and TARGATT™ provide justification for a more efficient and scalable system capable of differentiation into a variety of clinically-relevant tissue types^8–10^. While the Big-In system effectively demonstrated the use of thymidine-kinase as a counter-selective marker for large DNA insertions into landing pad H1-hiPSCs, the use of Cre recombinase as the core gene-transfer effector may lead to reduced accuracy and efficiency when compared to Bxb1^20^. The pluripotency and differentiation potential of the Big-IN system, however, was not validated. While STRAIGHT-IN utilizes Bxb1 and Cre recombinase for integration/excision respectively, the DICE and TARGATT systems, as well as other recombinase-based tools utilize the ϕC31 LSR as a core effector^26^. This enzyme exhibits substantial permissiveness for chromosomal rearrangements of the human genome and imprecise recombination in comparison to Bxb1^20,27,28^. As such, the use of RMCE catalyzed by Bxb1 alone may provide a safer, less off-target prone, transgene delivery to landing-pad hiPSCs. Furthermore, the use of a splice-acceptor gene trap in *AAVS1* driven by the endogenous *PPP1R12C* promoter can facilitate antibiotic resistance conditional on landing-pad targeting, enabling rapid isolation of recombined hiPSCs^29^.

Here we engineer the iPS11 human induced pluripotent stem cell line using CRISPR-HDR to insert landing-pad constructs positioned into the *AAVS1* genomic safe-harbour locus, and equip it with features which not only address the limitations of previous stem-cell-based landing-pad systems, but introduce novel features to expand the capabilities of these cells. Each construct is neomycin-resistant whilst expressing a red or green fluorescent protein marker. We conducted monoclonal expansion and downstream verification for each landing-pad construct, and quantified pluripotency and differentiability into all three germ-layers. Furthermore, both landing pad lines are homozygous, containing two sites for transgene integration. Herein we develop means of preferentially conducting heterozygous or homozygous transgenesis. As well, landing-pad hiPSCs are utilized as robust tools for synthetic biology by quantifying the recombination efficiency of all orthogonal Bxb1 *attP*/B site pairs, working towards targeted integration of multiple expression constructs in parallel. As such, we effectively equip investigators with a verified and flexible gene delivery toolkit capable of meeting the needs of cutting-edge of stem cell engineering.

## Results

### Landing-Pad Construct Design

Landing-pad HDR-donor constructs were generated by assembly of either EGFP, or mScarlet into an Ef1a-driven neomycin-resistance polycistron (Figure 1A). Constructs were flanked by Bxb1 *attB*(GT) and *attP*(GA) sites and homology arms for the *AAVS1* locus respectively. All constructs were verified by whole-plasmid sequencing to ensure functionality in downstream experiments.

**Figure 1:**
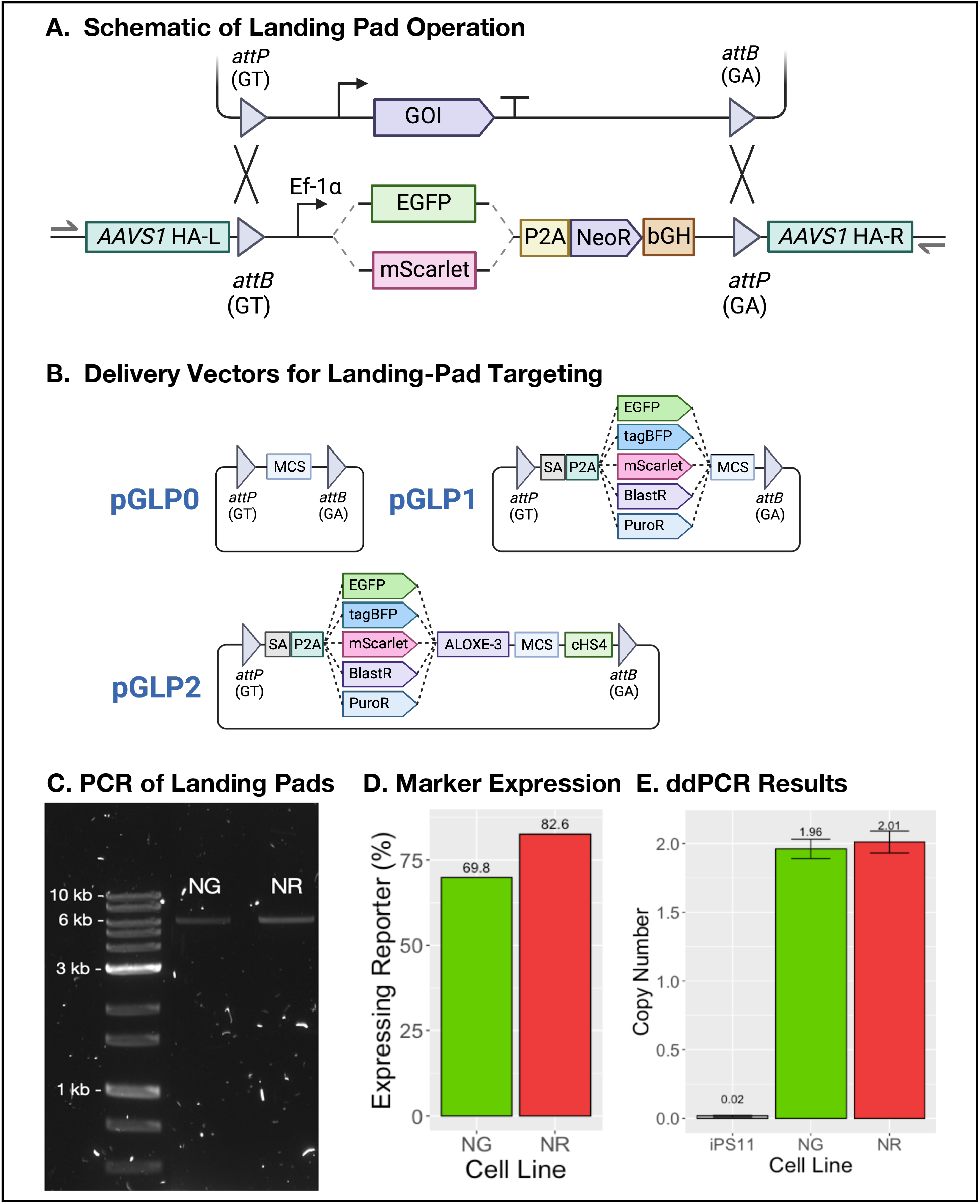
Landing-pad construct design and CRISPR-HDR knock-in. **(A)** Diagram of landing-pad construct design including Ef-1a-(EGFP/mScarlet)-P2A-NeoR expression cassette followed by bGH terminator sequence, orthogonal Bxb1 *attB* and *attP* recombination sites, and *AAVS1* homology arms (HAs). PCR primers for verification of landing-pad integration into the left and right sides of *AAVS1* are displayed as 1-sided arrows. A gene of interest flanked by compatible attP/B site pairs is depicted being targeted to the landing pad. **(B)** Schematic of delivery vector levels 0-3 are depicted with either puromycin and blasticidin antibiotic resistance markers or mScarlet, tagBFP or EGFP fluorescence markers. **(C)** Positive endpoint-PCR-based detection of landing-pad integration into *AAVS1*. **(D)** Reporter expression of EGFP in NG and mScarlet in NR cells following cell line generation **(E)** Copy-number of landing-pad constructs in the human genome.

### CRISPR HDR-Mediated Landing-pad hiPSC Integration and *AAVS1* Verification

Delivery of landing-pad DNA to hiPSCs followed by selection and clonal expansion yielded EGFP reporter landing-pad cells (classified as NG) and mScarlet reporters (classified as NR). Each were screened to ensure correct double-copy knock-in by PCR of the *AAVS1* locus with primers spanning regions distal to the repair template homology arms. Agarose-gel electrophoresis confirmed the expected sizes of 6661 bp for NG and 6640 bp for NR (Figure 1C). Nanopore sequencing of both NG and NR landing pad site PCRs followed by alignment with the CRISPR HDR repair-template plasmid confirmed correct insertion of all landing-pad components (Supplementary Information). After monoclonal expansion, NG and NR landing-pad cells were enriched by fluorescence-assisted cell-sorting (FACS), and flow-cytometric analysis conducted after cell-banking, thawing, and expansion revealed that 69.4% and 82.6% of NG and NR cells expressed their respective reporter gene (Figure 1D). Furthermore, absolute ddPCR quantification of Neomycin resistance gene copy compared to the human RNase P gene confirmed a double-copy of the landing pad within the genome (Figure 1E).

### Developing a Flexible Repertoire of Landing-Pad Delivery Vectors

In order to easily deliver a variety of gene constructs into landing-pad hiPSCs, a standard collection of vectors with sequentially increasing degrees of complexity were created (Figure 1B). The most basic “level-0” vector only contained a multiple-cloning site (MCS) flanked by *attP*/*attB*’ recombination sites. The “level-1” vectors incorporate splice-acceptor-based antibiotic resistance (puromycin or blasticidin) or fluorescent protein (EGFP, tagBFP, mScarlet) gene-trap cassettes. Additionally, flanking the MCS with ALOXE-3 and cHS4 transgene insulator sequences yielded the complete set of level-2 vectors^30^. All vectors were designed for compatibility with the MTK toolkit, creating an interface with extant modular cloning workflows^31^.

### Maintenance of Pluripotency, Baseline Differentiation Potential, and Karyotype in Landing-Pad hiPSCs

To ensure that genetic manipulation of the *AAVS1* locus and monoclonal expansion did not impact landing-pad hiPSC pluripotency, verification of *OCT-4* and *SOX-2* expression was determined (Figure 2A). Both NG and NR landing pad lines retained pluripotency marker expression in comparison to parental iPS11 cells. Differentiation of landing-pad cells into endoderm and ectoderm was also assessed in comparison to parental iPS11. For endodermal marker profiling, immunostaining and flow cytometric analysis of *CXCR4*, C-kit, and *SOX-17* expression after four days of differentiation revealed NG and NR cells exhibited double-positive populations similar to parental iPS11 cells (Figure 2B). Ectodermal *PAX-6*, maintenance of *SOX-2* and loss of OCT-4 profiling confirmed this finding (Figure 2C).

**Figure 2:**
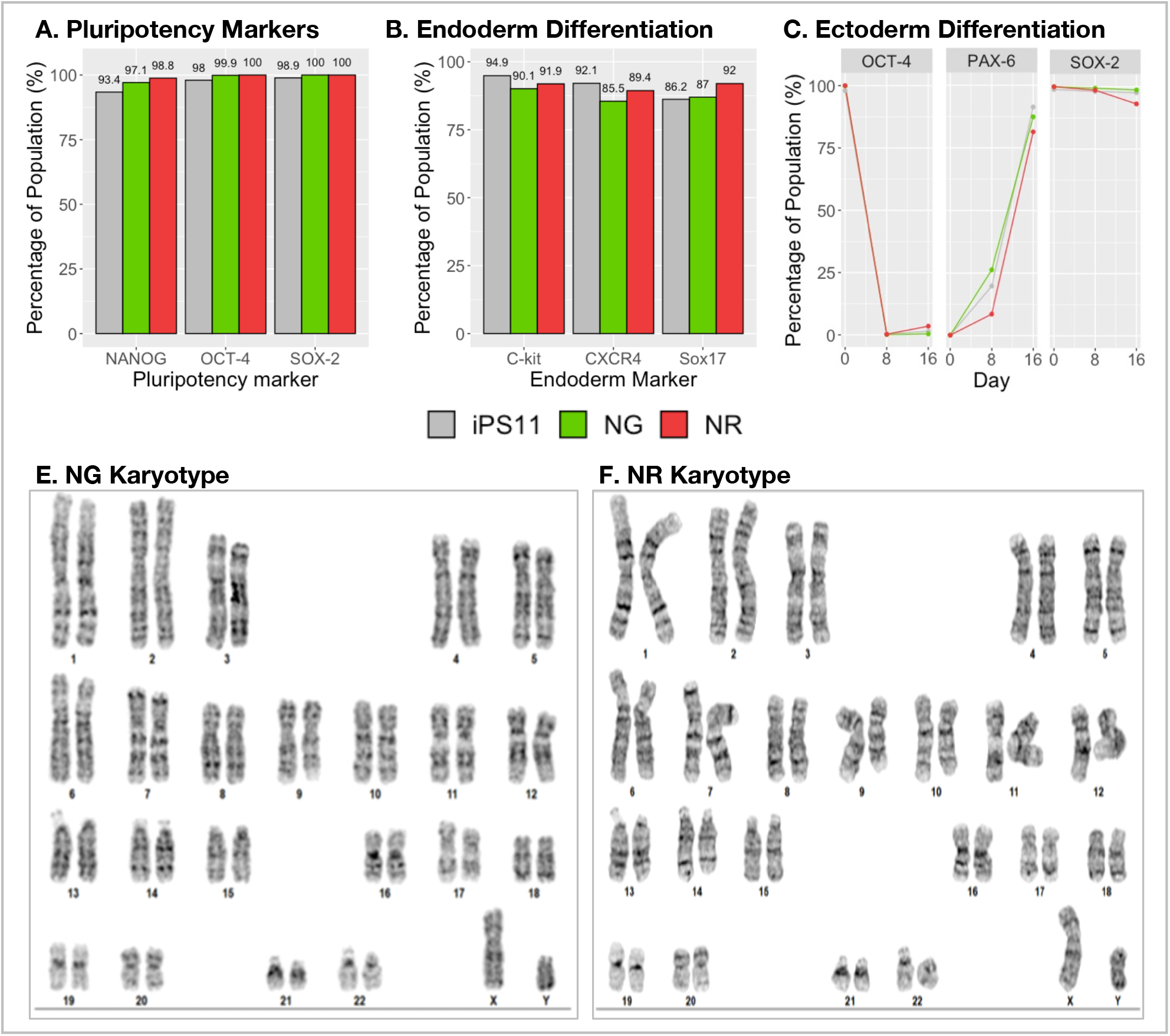
Pluripotency, differentiation marker, and karyotype profiling of NG and NR landing-pad hiPSCs. **(A)** Immunostaining and subsequent flow cytometric analysis of *OCT-4* and *SOX-2* expression markers compared landing-pad stemness to the parental hiPSC11 cell line. **(B)** Endoderm differentiation assessed by live-cell staining of C-kit and *CXCR4* expression as well as Sox17 expression in fixed cells in comparison to parental iPS11 cells. **(C)** Ectoderm differentiation assessed by loss of *OCT-4*, gain of *PAX-6*, and maintenance of *SOX-2* expression in landing-pad hiPSC sub-populations in comparison to parental iPS11 cells. **(D)** Karyotyping of NG and **(E)** NR Landing Pad hiPSCs. Representative G-banding for both cell lines are displayed. Both landing-pad hiPSCs maintain a 46X,Y karyotype.

Karyotyping of the NG and NR landing-pad hiPSC lines revealed that no chromosomal rearrangements had occurred during the CRISPR-HDR and monoclonal expansion processes (Figure 2D&E). Thus, these hiPSCs were deemed viable for testing of downstream applications.

### Functional Screening of Landing-Pad Integration Efficiency

To determine whether landing-pad cells could be effectively targeted for transgene delivery, flow cytometric quantification of an inserted fluorescent protein cassette was conducted. Due to the homozygosity of the landing-pad within the *AAVS1* locus, both dropout of the original landing-pad fluorescent marker to the delivered transgene, as well as the double-positive population were quantified for NG and NR landing-pad cells (Figure 3). For NG cells transfected with pGLP2-PuroR-EF1α-mCherry, It was determined that a mean 62.4% homozygous and 31.6% heterozygous mCherry-expressing population was isolated following 7-days of puromycin selection, whilst only 3.6% of cells did not receive a transgene. Similarly, NR cell transfected with pGLP2-Puror-Ef1α-EGFP exhibited a mean 75.1% homozygous and 20.9% heterozygous EGFP-expressing population, whilst only 2.2% of cells did not receive a transgene.

**Figure 3:**
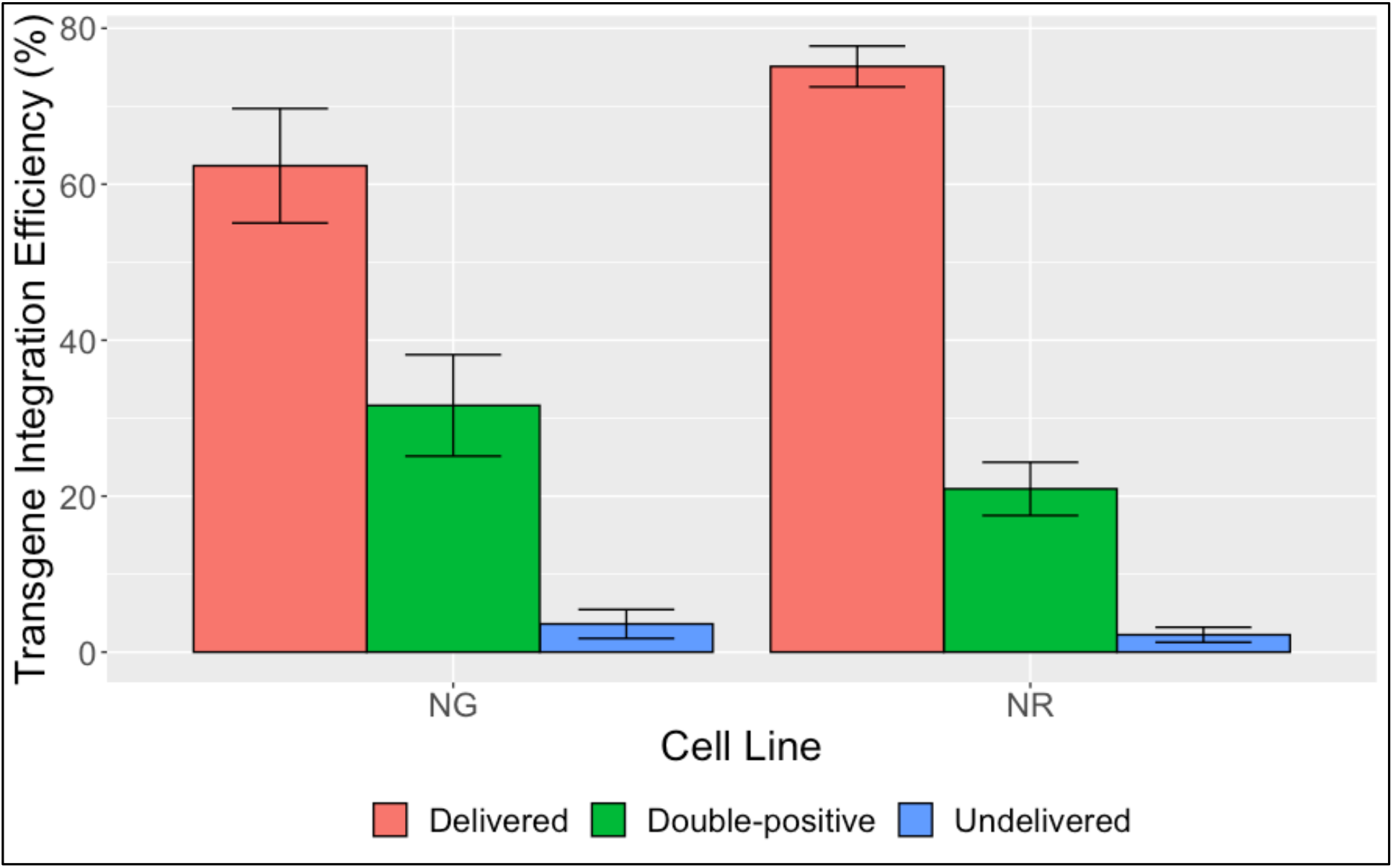
Functional screening of landing-pad integration efficiency. Ef1a-mCherry and Ef1a-EGFP transgenes were delivered to NG and NR landing-pad hiPSCs respectively followed by 7-days of puromycin selection (0.25 ug/mL). Error bars represent standard deviation of triplicate samples.

### Confirmation of Landing Pad hiPSC Differentiation Potential After Transgene Delivery

Following transfection, antibiotic-enrichment, and FACS for selection of single and double-positive reporter-expressing cell populations, we reattempted differentiation into endoderm and ectoderm to confirm that manipulation of the landing-pad with Bxb1 did not impact their pluripotency. We confirmed that NG and NR heterozygotes (1C) and homozygotes (2C) retained differentiation potential (Figure 4A-B).

**Figure 4:**
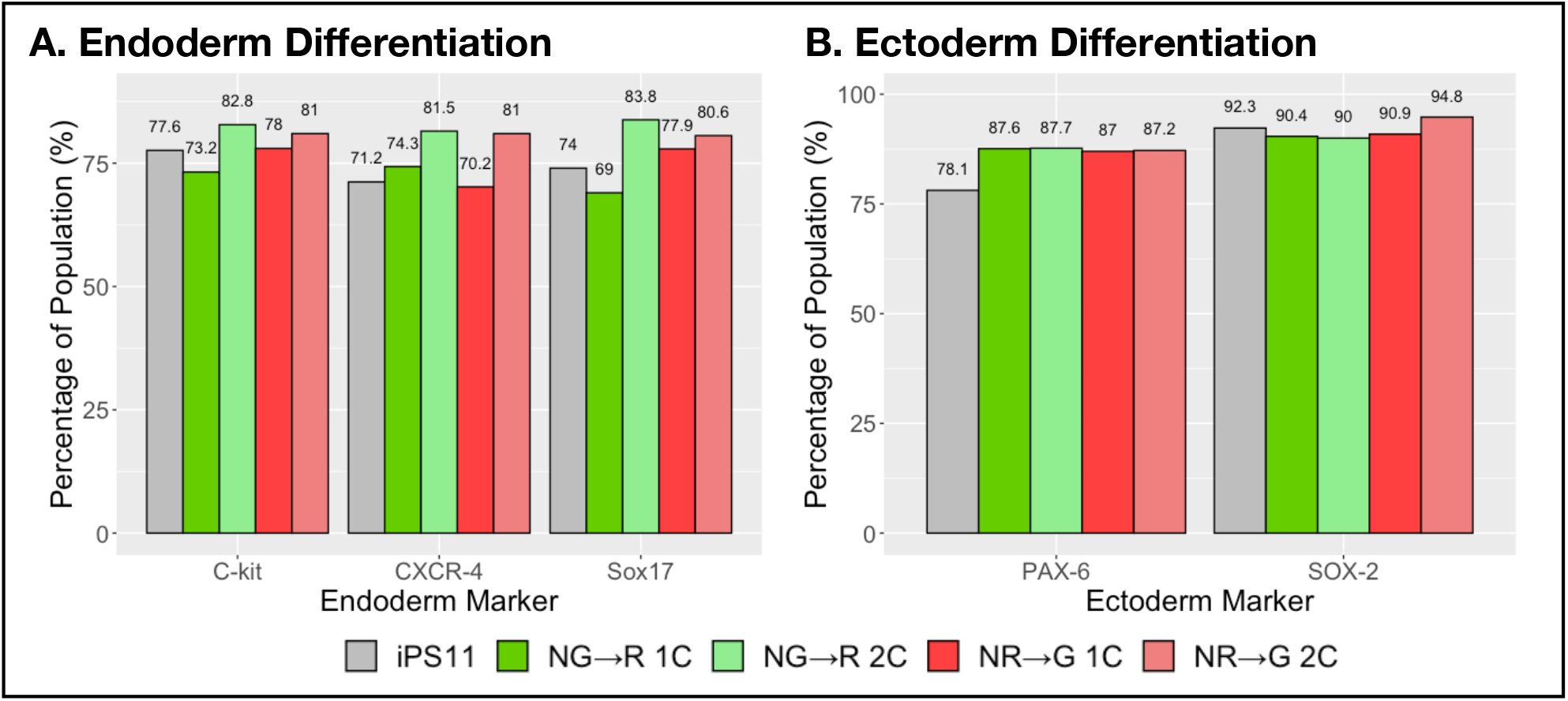
Differentiation marker profiling of NG and NR landing-pad hiPSCs with loaded transgenes. Analysis of parental (iPS11) as well as NG cells transfected with mCherry transcriptional unit with double-positive heterozygous (NG→ R 1C) or fully-delivered homozygous (NG→ R 2C) configuration, or NR cells transfected with EGFP transcriptional unit with double-positive heterozygous (NR→ G 1C) or fully-delivered homozygous (NR→ G 2C) configuration. **(A)** Endoderm differentiation assessed by live-cell staining of C-kit and *CXCR4* expression as well as Sox17 expression in fixed cells in comparison to parental iPS11 cells. **(B)** Ectoderm differentiation assessed by *PAX-*6 and SOX*-2* expression in landing-pad hiPSC sub-populations in comparison to parental iPS11 cells assessed on day 16 of neural differentiation.

### Differentiation of Landing Pad hiPSCs into Terminal Lineages

We next aimed to confirm that the landing-pad hiPSC lines were competent to be directed to specific cell lineages with the same robustness as the iPS11 parental line. NR landing-pad hiPSCs were directed to the skeletal muscle myogenic progenitor lineage with a similar frequency as observed with the iPS11 cells (Figure 5A) and produced phenotypically similar cultures of multinucleated, striated skeletal muscle myotubes (Figure 5B-C) when placed in low serum conditions.

**Figure 5:**
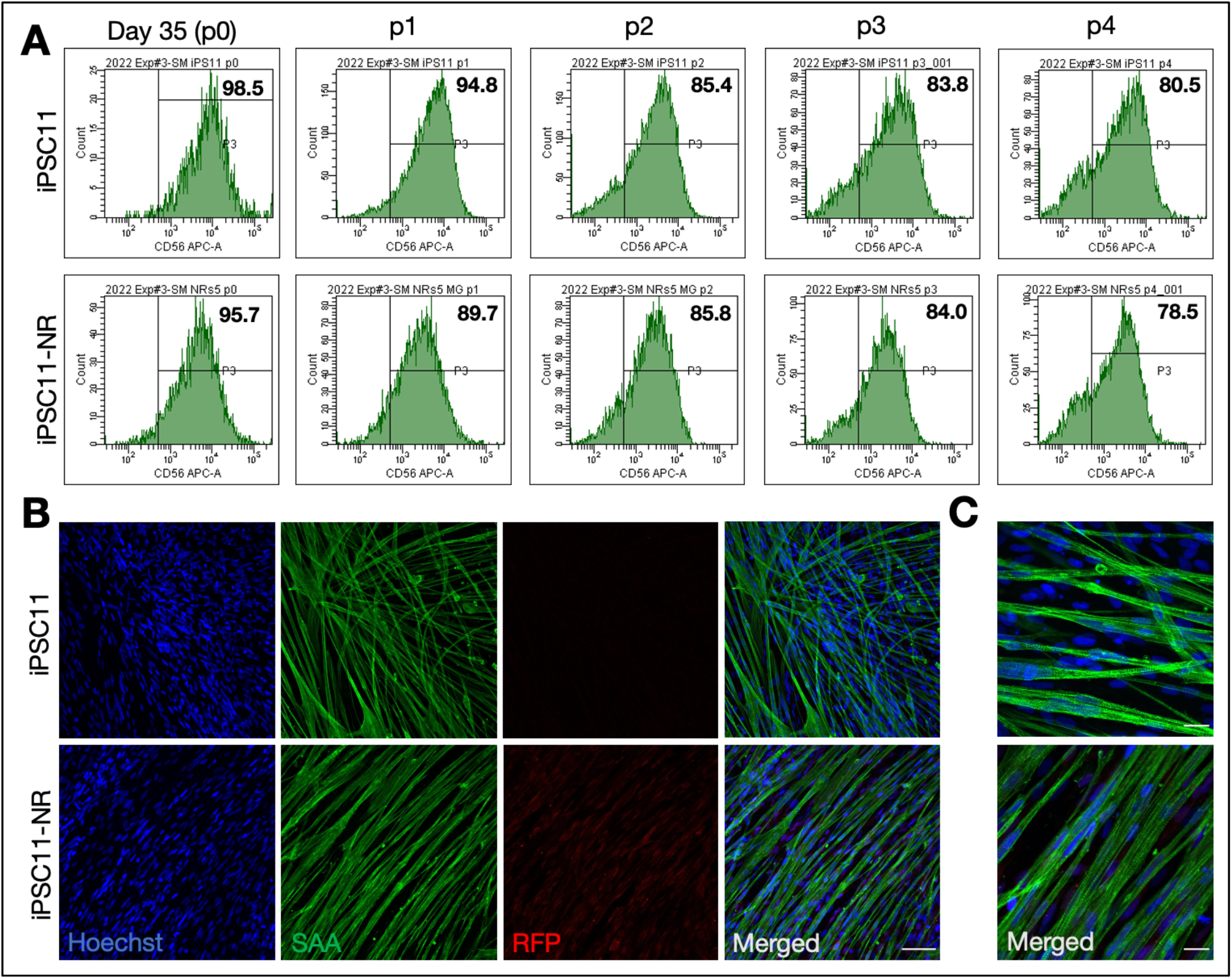
NR landing-pad hiPSC direction to skeletal muscle cells. **(A)** Flow cytometric graphs of CD56 expression in populations of iPS11 and iPS11-NR derived myogenic progenitors across passages 0 – 4 cultures. Schematic of hiPSC direction to myogenic progenitors and subsequent differentiation to multinucleated myotubes. **(B-C)** Representative **(B)** 20x and **(C)** 60x confocal images of multinucleated skeletal muscle cells produced from (top) iPS11 and (bottom) NR landing-pad iPSCs and immunostained to visualize nuclei (Hoechst, blue), sarcomeric alpha-actinin (SAA, green), and red fluorescent protein (RFP, red). Scale bars (C) 25 um and (D) 100 um.

## Methods

### Landing-Pad Donor and Cas9/gRNA Expression Construct Assembly

In order to construct each donor vector, the backbone of pSH-EFIRES-P-AtAFB2-mCherry (Addgene #129716) was PCR-amplified to include both homology arms to *AAVS1*. Both Bxb1 *attP*(GA) and *attB*(GT) sites were synthesized by oligonucleotide overlap assembly as designed by *Primerize*^32^. Other fragments encoding the Ef1a promoter, EGFP, mScarlet, P2A, and neomycin resistance were PCR-amplified from extant constructs present in our laboratory using NEB *Q5* Hot-Start 2X Mastermix according to the manufacturer’s protocol (New England Biolabs, Ipswitch, MA, USA; M0494L). HDR-donor assemblies were completed using *NEBuilder HiFi 2X Mastermix* in accordance with the manufacturer’s protocol (New England Biolabs; E2621L). HDR-donor constructs were verified by whole-plasmid sequencing by Primordium Labs (Arcadia, CA, USA). An all-in-one Cas9/gRNA expression construct by cloning an *AAVS1* sgRNA sequence derived from pCas9-sgAAVS1-2 PX458 (Addgene #129727) downstream of the hU6 promoter in a PX458 (Addgene #48138) Cas9 expression vector by golden-gate assembly with BbsI-HF (New England Biolabs, R3539L)^33,34^. All plasmids were transformed into NEB *Stable E. coli* and grown at 30°C to ensure construct stability (New England Biolabs). Plasmid clones were picked and inoculated into liquid LB media for overnight culture followed by miniprep using the PureLink Plasmid MiniPrep Kit (Invitrogen, Waltham, MA, USA; K210010), and verified by sanger-sequencing by Eurofins genomics (Louisville, KY, USA). Correct plasmid clones were further expanded in 100 mL overnight cultures in terrific broth, and isolated using the ZymoPure II Plasmid MaxiPrep Kit (Zymo Research, Irvine, CA, USA; D4203).

### Long single-stranded DNA (lssDNA) HDR-Donor Generation

The lssDNA generation protocol was adapted from Rosenstein and Walker^35^. Briefly, PCR amplification of each donor construct and flanking *AAVS1* homology arms was conducted using a forward primer modified with a 5’-biotin moiety and five 5’-phosphorothioate linkages, and a 5’-phosphorylated reverse primer. 200 μL reactions using Herculase II DNA polymerase were conducted according to the manufacturer’s protocols (Agilent, Santa Clara, CA, USA; 600675). PCR reactions were purified and digested with 10U of lambda exonuclease (New England Biolabs; M0262) in 1X lambda-exonuclease buffer for 2 hours at 37°C, followed by addition of 10 U dsDNAse (Thermofisher Scientific, Waltham, MA, USA; EN0771) for 10 min, and heat inactivation at 65°C for 20 min. Reactions were purified according to the ssDNA cleanup protocol from the Monarch PCR cleanup kit (New England Biolabs; T1030L).

### hiPSC Culture

iPS11 hiPSCs (Alstem Cell Advancements, Richmond, CA, USA; iPS11) cultured with mTeSR-Plus (StemCell Technologies, Vancouver, BC, Canada; 100-0275) according to the manufacturer’s protocol on GelTrex-coated plates (Invitrogen; A1413201). RevitaCell supplement was added at 1X concentration for all seeding and passaging steps (Invitrogen; A2644501). Cells were grown in a 5% CO_2_ incubator set to 37°C. Passaging was conducted using TrypLE gentle-dissociation reagent according to standard tissue culture protocols (Invitrogen; 12604013).

### CRISPR-HDR Mediated Knock-In of Landing-Pad Donor DNA in hiPSCs

hiPSCs were grown to 60% confluence in a 24-well plate, and media was changed and supplemented with 1X RevitaCell 30 min prior to transfection with a 500 ng of a 1:1 mixture of PX458:lssDNA donor for each landing-pad construct using lipofectamine stem in accordance with the manufacturers protocol (Invitrogen; STEM00001).

### Selection and Clonal Expansion of Landing-Pad hiPSCs

24-hours after transfection, cells from each well were dissociated and resuspended in cloning media consisting of mTeSR-Plus supplemented with 1X CloneR2 (StemCell Technologies; 100-0691) and mixed gently with a P1000 pipette tip in order to generate a single-cell suspension. Cell samples were counted using a Countess II Haemocytometer (Life Technologies, Carlsbad, CA, USA), and plated onto 10 cm GelTrex-coated tissue culture plates at a density of 50 cells/cm^2^ with 8 mL of cloning media. 2 days later, culture media was changed with fresh cloning media. On the 4^th^ day, cloning media was removed and replaced with mTeSR-Plus supplemented with 50 μg/mL G418 selective antibiotic. For the next ten days, media changed every other day with increasing concentrations of G418 (Invitrogen; 10131035) up to 250 μg/mL. Once discrete 200-300 cell colonies for each dish were visible, eight colonies per-plate were picked, dissociated, expanded, and split into duplicate 24-well plates.

### Fluorescence-Activated Cell Sorting (FACS) of hiPSCs

hiPSCs were grown to 80% confluency in 6-well plates and were dissociated with 750 μL of TrypLE. Following incubation at 37°C for 5 mins, 750 μL of mTeSR-plus was added, and cells were pelleted by centrifugation at 200xg for 5 mins. Cells were resuspended in sorting buffer (DPBS supplemented with 5 mM EDTA and 1% BSA) at a density of 6 million cells/mL, and were pipette-mixed vigorously to dissociate cell aggregates. Cells were filtered through a 40 μm nylon cell-strainer into a 5 mL polystyrene FACS tube. A collection tube of 2 ml mTeSR-plus supplemented with 1X RevitaCell was prepared for deposition of sorted cells. Cells were sorted on a FACS Aria IIIu by staff members of the University of Toronto -Temerty Faculty of Medicine Flow Cytometry Facility staff members. NG cells were gated on +EGFP expression, while NR cells were gated on +mScarlet expression. Cells were cultured for 4-days undisturbed post-sorting before downstream manipulation.

### Verification of Landing-pad Insertion

While one of the 24-well plates containing all landing-pad hiPSC clones was maintained for further growth, the second was used for gDNA extraction. gDNA was extracted using the Invitrogen PureLink Genomic DNA Mini Kit. PCR was conducted using primers flanking the *AAVS1* homology arms (AAVS1_screenF: ACCTGCCCAGTACAGGCATC, AAVS1_screenR: TCCCCGTTGCCAGTCTCGAT) present on the landing-pad donor construct. 100 ng of gDNA was mixed with AAVS1_screenF/R primers to a final concentration of 500 nM, as well as 3% v/v DMSO and 50% with Phire II Hot-Start PCR 2X mastermix (Thermo Scientific; F125L), and topped-up to 25 μL with ddH2O. PCR was run at an initial denaturation temperature of 98°C for 30 sec, followed by 35 cycles of 98°C for 15 sec, 68°C for 20 sec, and 72°C for 2:45 min, and finished by a 2 min final extension at 72°C. PCR products were purified with the Monarch PCR cleanup kit according to the manufacturer’s protocol. Agarose gel electrophoresis of each PCR product was conducted to confirm correct homozygous integration of landing-pads into the *AAVS1* site. ddPCR was conducted by the Center for Applied Genomics (TCAG, Toronto, ON, Canada) in order to calculate the copy-number of landing-pad constructs in each candidate cell line by absolute quantification of the neomycin resistance gene (present in on landing-pad constructs) normalized to the human RNase P gene.

### Flow Cytometry of hiPSC Pluripotency Markers

Staining for *OCT-4, SOX-2, and NANOG* was conducted with the following primary antibodies respectively: anti-Oct-3/4 (BD Biosciences, San Jose, CA, USA), rabbit anti-Sox-2 (New England Biolabs), and rabbit anti-NANOG (New England Biolabs). Briefly, cells were detatched with TrypLE (Invitrogen) and quenched with mTeSR Plus, followed by a 5 min centrifugation at 200xG for 5 minutes, followed by a wash-step with PBS. Cells were then incubated with respective primary antibodies at room temperature for 1 hour, followed by two PBS-wash steps and secondary antibody staining and analysis on an LSRFortessa X-20 flow cytometer (BD Biosciences), which was also used for all downstream flow cytometry.

### Differentiation of hiPSC to Endoderm and Ectoderm and Confirmation Thereof

Endoderm differentiation was adapted from the stage-1 protocol described in Nostro *et. al*., with cells maintained in d1-media until day-4 of differentiation^36^. Cells were then dethatched, PBS-washed, and split for subsequent endodermal marker staining. A third of each cell sample was stained for 30 mins at 4°C with 1:100-dilutions of APC-conjugated anti-*C-Kit*/CD117 (BD Biosciences; 550412) and PE-Cy7-conjugated anti-*CXCR-4*/CD184 (BD Biosciences; 560669), followed by two washes with PBS+5% FBS. The remaining sample was fixed with 4% paraformaldehyde for 10 mins on ice, followed by permeabilization with ice-cold absolute methanol for 3 mins and washing with PBS+5% FBS. Fixed cells were stained with 1:100 dilution of APC-conjugated anti-*SOX-17* (R&D Systems, Minneapolis, MN, USA; IC1924A) for 30 mins at 4°C, followed by two washes with PBS+5% FBS and flow cytometric analysis.

Ectoderm differentiation was conducted using the STEMdiff™ SMADi Neural Induction Kit (Stemcell technologies; 08581) according to the manufacturer’s protocol. On day-8 and/or 16 of differentiation, cells were detatched, PBS-washed, and fixed with 4% PFA for 15 mins on ice, followed by permeabilization with ice-cold absolute methanol for 3 mins. Cells were then washed with PBS+5% FBS and stained with 1:100-dilutions of AF647-conjugated anti-PAX-6 (BD Biosciences; 562249) and PerCP-Cy5.5-conjugated anti-SOX-2 (BD Biosciences; 561506).

### Functional Screening of Landing-Pad Integration Efficiency

In order to quantify landing pad hiPSC recombination efficiency, the *NG* and *NR* cell lines were transiently co-transfected with pEf1a-Bxb1-NLS and pGLP2-based delivery vector (pGLP2-Ef1a-mCherry for NG, pGLP2-Ef1a-EGFP for NR) with Lipofectamine stem in a 3:1 plasmid mass ratio. As well, control reactions excluding the Bxb1-integrase expression vector were ran to determine when transient transfection of delivery vector had subsided. ∼48 hrs post-transfection, 0.25 ug/mL puromycin selective antibiotic (Thermofisher Scientific) was added to culture, and media was changed daily until no live-cells were visible in the no-integrase control well. Cells were dissociated with TrypLE, and replated for further expansion and downstream FACS.

### Assembly of Landing-Pad Delivery Vectors

Plasmid DNA assembly of landing pad constructs was completed using *NEBuilder HiFi 2X Mastermix* in accordance with the manufacturer’s protocol (New England Biolabs), as well as *Primerize* for *attB* and *attP* site assembly. Delivery vectors were verified by whole-plasmid sequencing by Primordium Labs^32^.

### Differentiation of hiPSC to Myogenic Progenitors and Skeletal Myotube

hiPSCs were differentiated to the myogenic lineage to produce skeletal muscle myogenic progenitors by following the detailed protocol established by Xi *et. al*.^37^. Minor modifications to some of the reagents used are detailed here: Geltrex (ThermoFisher Scientific; A1413302; 2 %) rather than Matrigel, addition to DNase I (10 ug/ml: Millipore Sigma, Burlington, MA, USA; 260913) to collagenase IV (ThermoFisher Scientific; 17104019) and TrypLE during the day 29 dissociation step, and SK Max medium (Wisent Bioproducts, Saint-Jean Baptise, QC, CA; 301-061-CL) as an alternative to SkGM2. Upon establishing a myogenic progenitor line, a portion of passage 0 cells were cryopreserved for future studies. Myogenic progenitors were used for experiments at passages 1 – 3.

To generate multinucleated myotube cultures, iPS11 and iPS11-NR derived myogenic progenitors were seeded on Geltrex coated (1:100) plates in Wisent SK Max medium supplemented with 10 % FBS and 20 ng/mL FGF2. Upon reaching 70-80% confluence the media was switched to a differentiation inducing medium consisting of DMEM with 2 % horse serum and bovine insulin (10 ug/mL) for 4-6 days. The differentiation media was exchanged every two days. Myotube cultures were fixed, stained, and imaged exactly as previously described^38^.

## Discussion

The core development of this study is the design and verification of a Bxb1-based RMCE landing-pad placed into the *AAVS1* safe-harbour locus using CRISPR-HDR in hiPSCs. While both plasmid and lssDNA-based HDR donors were utilized to knock-in cells, this study presents the first lssDNA-based donor strategy in hiPSCs using chemical transfection, which confirms previous reports of this method outperforming plasmid-based CRISPR-HDR in other systems^39^. We expect this method will be an effective tool for stable transfection without access to a nucleofector, simplifying cell-line generation and potentially reducing associated cytotoxicity during transfection.

An in-depth slate of experiments for verification of landing-pad hiPSC validity was conducted (Figures 1,2) to ensure that the cells developed herein would be capable of downstream use. PCR of the *AAVS1* locus containing the landing pad, followed by nanopore-sequencing leveraged a recent trend in cost-effective whole-plasmid and PCR product sequencing. We expect that investigators may use similar methodologies to confirm transgene delivery with sequence-level certainty. While our initial experimentation on landing pad cells placed an emphasis on single-copy landing-pad gene integration, we later decided that homozygous landing-pad lines presented investigators with increased flexibility for transgenesis, and as such the decision was made to preferentially screen for homozygous hiPSC clones. While the percentage of landing pad cells expressing the fluorescent reporter declined even post-sort to ∼70-80% (Figure 1D), ddPCR conducted from gDNA templates isolated from the same cells confirmed homozygosity of both landing-pad lines, indicating that transgene silencing may be the cause of this decline as opposed to loss of the landing-pad from the genome.

Confirmation of pluripotency of both NG and NR hiPSC clones (Figure 2A), as well as differentiation into endoderm and ectoderm (Figure 2B,C) was essential to validating the utility of the landing-pad cells for general use by investigators with a wide range of specific cell-types in-mind. Confirmation of the normal karyotype (Figure 2D,E) of both lines was the final indication that genetic manipulation of these cells did not adversely effect their genomic integrity, and as such act as a faithful human stem cell model for downstream use. Furthermore, integration of transgenes at the landing pad did not impact downstream differentiation potential to endoderm and ectoderm (Figure 4), which fully validates the genetic engineering arc of these cells from generation to end-use. While mesodermal differentiation was not directly assessed, validation of skeletal muscle differentiation as depicted in Figure 5 confirms that the cells generated herein are capable of being directed towards this lineage.

During functional screening of the landing pad cells to confirm transgene delivery as depicted in Figure 3, it was observed that a mixed population of either homozygotes or heterozygotes existed for both NG and NR cells. Interestingly, homozygosity was favourable over heterozygosity. Yet, considering that two independent Bxb1-mediated integration events (two sets of attP/attB pairs) are required for full transgene insertion into the landing-pad, one might surmise that the probability of their intersection would be lower. It may be the case that cells which receive a large-enough dose of Bxb1 to undergo a single-integration event are poised to continue landing-pad recombination to completion, and as such heterozygosity would be unfavourable. Regardless, the relative abundance of either transgene configuration was deemed an acceptable result, as whether an investigator desired homozygotes or heterozygotes, sorting for ∼70% or %30 of a cell population respectively is a generally attainable task.

It is our sincere hope that the landing pad lines described herein will be an accessible tool for the stem-cell community to rapidly engineer custom cells for downstream use. Whether for generating stable genomically-integrated complex gene circuit schemes for the synthetic-biology, or copy-number related disease modelling, this system can accommodate a wide variety of potential experiments. Furthermore, it may be possible to simplify library-screening efforts in stem cells or downstream terminal lineages with this landing pad system by enriching for single-copy transgene integration events representative of large variant libraries. Regardless of purpose, we believe our contribution to furthering the utility of landing-pads in hiPSCs is well-warranted and can accommodate the future of stem-cell engineering and regenerative medicine.

## Conclusion

This report aimed at characterizing the use of landing-pads for robust engineering of hiPSCs, hopefully enabling more facile and customizable transgenesis and powerful library screening of clinically relevant cell types. We were able to successfully engineer the iPS11 cell line with landing-pad constructs expressing one of a green or red fluorescent marker, and confirm their suitability for use in downstream experiments by pluripotency marker analysis, differentiation potential testing, and functional screening. Furthermore, we designed a flexible repertoire of entry vectors for this system, rapidly generating fully-selected stably-expressing cell line by landing-pad gene delivery. As such, we believe that the landing-pad hiPSC toolkit will be of invariable benefit for the stem cell research and regenerative medicine community for the years to come.

